# Retinoblastoma 1 (*RB1*) modulates the proliferation of chicken preadipocytes

**DOI:** 10.1101/341453

**Authors:** Yu-Xiang Wang, Hai-Xia Wang, Wei Na, Fei-Yue Qin, Zhi-Wei Zhang, Jia-Qiang Dong, Hui Li, Hui Zhang

## Abstract

Retinoblastoma 1 (*RB1*) has been extensively studied in mammalian species, but its function in avian species is unclear. The objective of this study was to reveal the role of chicken *RB1* (Gallus gallus *RB1*, *gRB1*) in the proliferation of preadipocytes. In the current study, quantitative real-time PCR analysis showed that the expression levels of *gRB1* transiently increased during the proliferation of preadipocytes. The MTT assay showed that *gRB1* overexpression suppressed preadipocyte proliferation, and *gRB1* interference promoted preadipocyte proliferation. Additionally, cell-cycle analysis indicated that *gRB1* may play a crucial role in the G1/S transition. Consistently, gene expression analysis showed that *gRB1* knockdown promoted marker of proliferation Ki-67 (*MKi67*) expression at 96 h (*P* < 0.05), and that overexpression of *gRB1* reduced *MKi67* expression at 72 h (*P* < 0.05). Together, our study demonstrated that *gRB1* inhibited preadipocyte proliferation at least in part by inhibiting the G1 to S phase transition.

## Introduction

The selection for rapid growth in meat-type chickens has been accompanied by increased abdominal fat deposition [1]. Excessive abdominal fat deposition can decrease feed efficiency and carcass quality, leading to consumer rejection of the meat [2-4]. The excessive deposition of abdominal fat is mainly due to the excessive proliferation and differentiation of adipocytes in adipose tissues. Clarifying the genetic mechanisms of proliferation and differentiation of adipocytes will help control the excessive accumulation of abdominal fat.

In our previous study, we mapped a quantitative trait locus (QTL) with major effects on abdominal fat trait into a 3.7-Mbp (172.6–176.3 Mbp) region in chicken chromosome 1 using an F2 population of a broiler × layer cross [5]. In this 3.7-Mbp region, five genes, including Retinoblastoma 1 (*RB1*), were detected [6]. In the examination of genome-wide selection signatures of abdominal fat content with the chicken 60 k SNP chip and the extended haplotype homozygosity (EHH) assay, a number of genes in the significant core regions were detected, and the *RB1* gene was detected once again [7]. In addition, polymorphism analysis of the *RB1* gene of Northeast Agricultural University broiler lines divergently selected for abdominal fat content (NEAUHLF) indicated that the polymorphisms of *RB1* were associated with abdominal fat content (unpublished data). These results suggest that *RB1* plays an important role in the deposition of abdominal fat in broiler chickens. However, the mode of action of *RB1* in chicken abdominal fat deposition remains unknown, hence, the aim of this study was to analyze the function of *RB1* in the proliferation of chicken preadipocytes.

## Materials and methods

### Ethics statement

All animal work was conducted according to the guidelines for the Care and Use of Experimental Animals established by the Ministry of Science and Technology of the People’s Republic of China (approval number: 2006–398), and was approved by the Laboratory Animal Management Committee of Northeast Agricultural University.

### Preparation and culture of cells

In the current study, the immortalized chicken preadipocyte line (ICP1) was used to analyze the function of *RB1* in the proliferation of chicken preadipocytes. Primary chicken preadipocytes were isolated from the abdominal adipose tissue of 10-day-old Arbor Acres (AA) broilers, and then the primary chicken preadipocytes were infected with either chicken telomerase reverse transcriptase (chTERT) alone or in combination with chicken telomerase RNA (chTR) to establish immortalized chicken preadipocyte line (ICP1) [8]. ICP1 had survived >100 population doublings *in vitro* and displayed high telomerase activity and had no sign of replicative senescence [8]. This cell line shows great promise as an *in vitro* model for the investigation of chicken adipogenesis and lipid metabolism. ICP1 cells were maintained in DMEM F12 supplemented with 10% fetal serum and 1% penicillin and streptomycin. ICP1 cells were cultured until 90% confluence and then passaged and seeded in cell culture plates at a density of 1 × 10^5^ cells/cm^2^.

### Construction of RB1-overexpression plasmid and synthesis of siRNA-RB1

To carry out the overexpression and RNA interference (RNAi) experiments, the *RB1*-overexpression plasmid was constructed and siRNA-*RB1* was synthesized. The full-length coding sequence of chicken *RB1* (Gallus gallus *RB1*, *gRB1*; GenBank accession number: NM_204419) was amplified from chicken abdominal adipose tissue cDNA using a pair of specific primers: sense, 5′-ACGTCGACAACGGTCACCATGCCGCCC-3′; anti-sense, 5′-CCGCTCGAGCAGCCCCTGGTCCTGAGGAGAATC-3′. The PCR product was cloned into pEasy-T1 Simple vector (TransGen, Beijing, China) and verified by direct sequencing. The full-length coding sequence of *gRB1* was excised from the pEasy-T1-*gRB1* plasmid by digesting with SalI and XhoI, and subcloned into pCMV-HA vector (Clontech, Mountain View, CA, USA) to obtain the *RB1*-overexpression vector, pCMV-HA-*gRB1*.

The siRNA of the *gRB1* selected for RNAi and the negative control were designed and synthesized by GenePharma Company (Shanghai, China; Table 1).

**Table 1.**
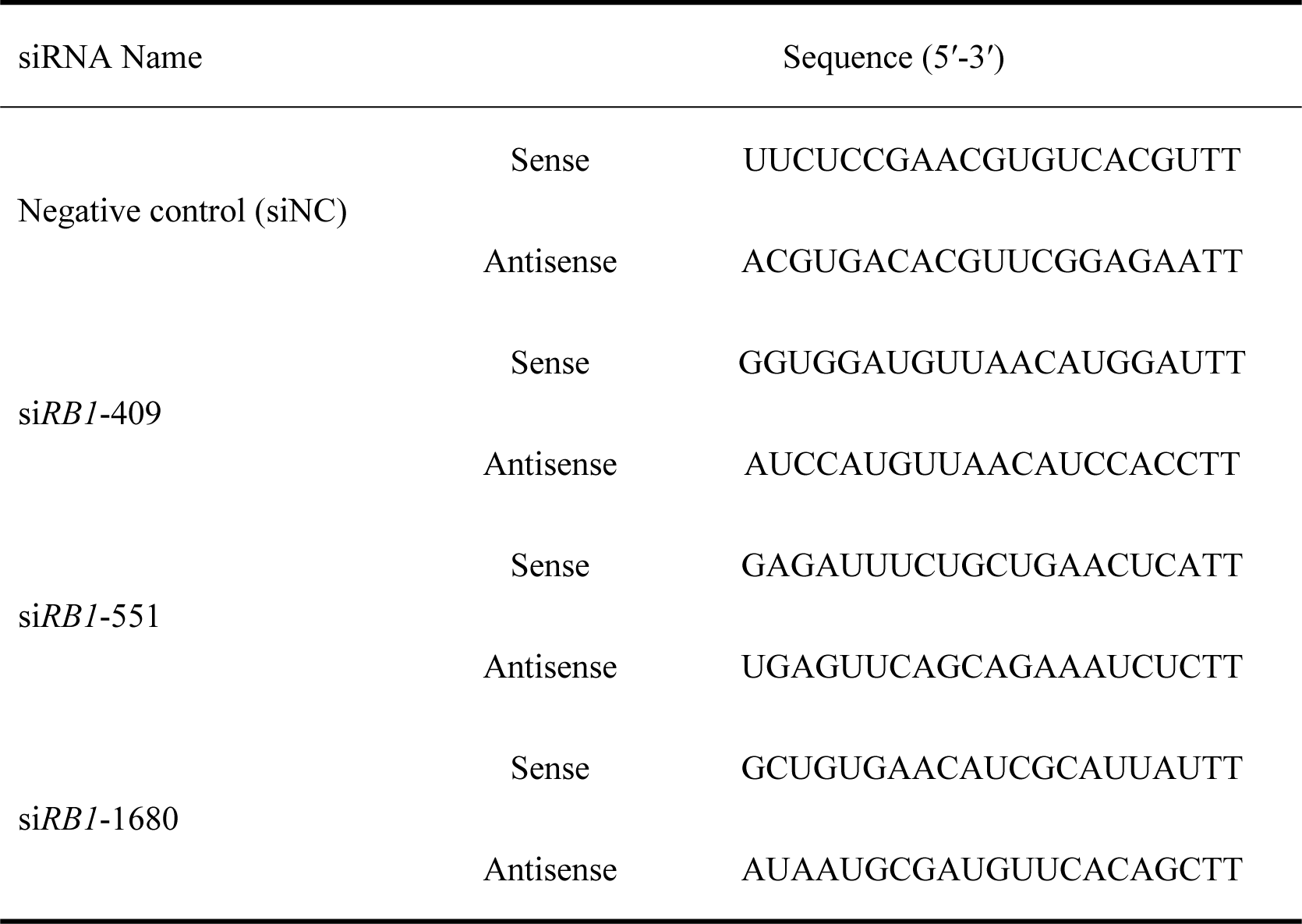
siRNA Sequences.

### RNA Isolation and real-time RT-PCR

To detect the expression level of *RB1*, marker of proliferation Ki-67 (*MKi67*), *Cyclin D1*, proliferating cell nuclear antigen (*PCNA*) and Transcription factor E2F1 (*E2F1*) genes in preadipocytes, the RNA of the cells was isolated and real-time RT-PCR was carried out. Total RNA was isolated using Trizol reagent (Invitron, Monmouth, UK). RNA quality was assessed by denaturing formaldehyde agarose gel electrophoresis. Reverse transcription was performed using 1 μg of total RNA, an oligo (dT) anchor primer, and EasyScript reverse transcriptase (TransGen). Reverse transcription conditions for each cDNA amplification were 42°C for 30 min, and then 85°C for 5 min.

Real-time RT-PCR was carried out using the 7500 real-Time PCR System (Applied Biosystems, Foster City, CA, USA) and the TransStart Top Green qPCR SuperMix (TransGen). The primers for *RB1*, *MKi67*, *Cyclin D1*, *PCNA*, and *E2F1* genes used for real-time RT-PCR are shown in Table 2. Non-POU domain-containing octamer-binding (*NONO*) was used as the internal reference gene. Dissociation curves were analyzed using the Dissociation Curve 1.0 software (Applied Biosystems) for each PCR reaction to detect and eliminate possible primer-dimer artifacts. Results (fold changes) are expressed as 2^-ΔCt^ in which ΔCt = (Ctij − Ctrj), where Ctij and Ctrj are the Ct for gene i and reference gene r in the sample (named j). The statistical significance of the differences in mRNA expression levels between groups was determined by the *t*-test.

**Table 2.**
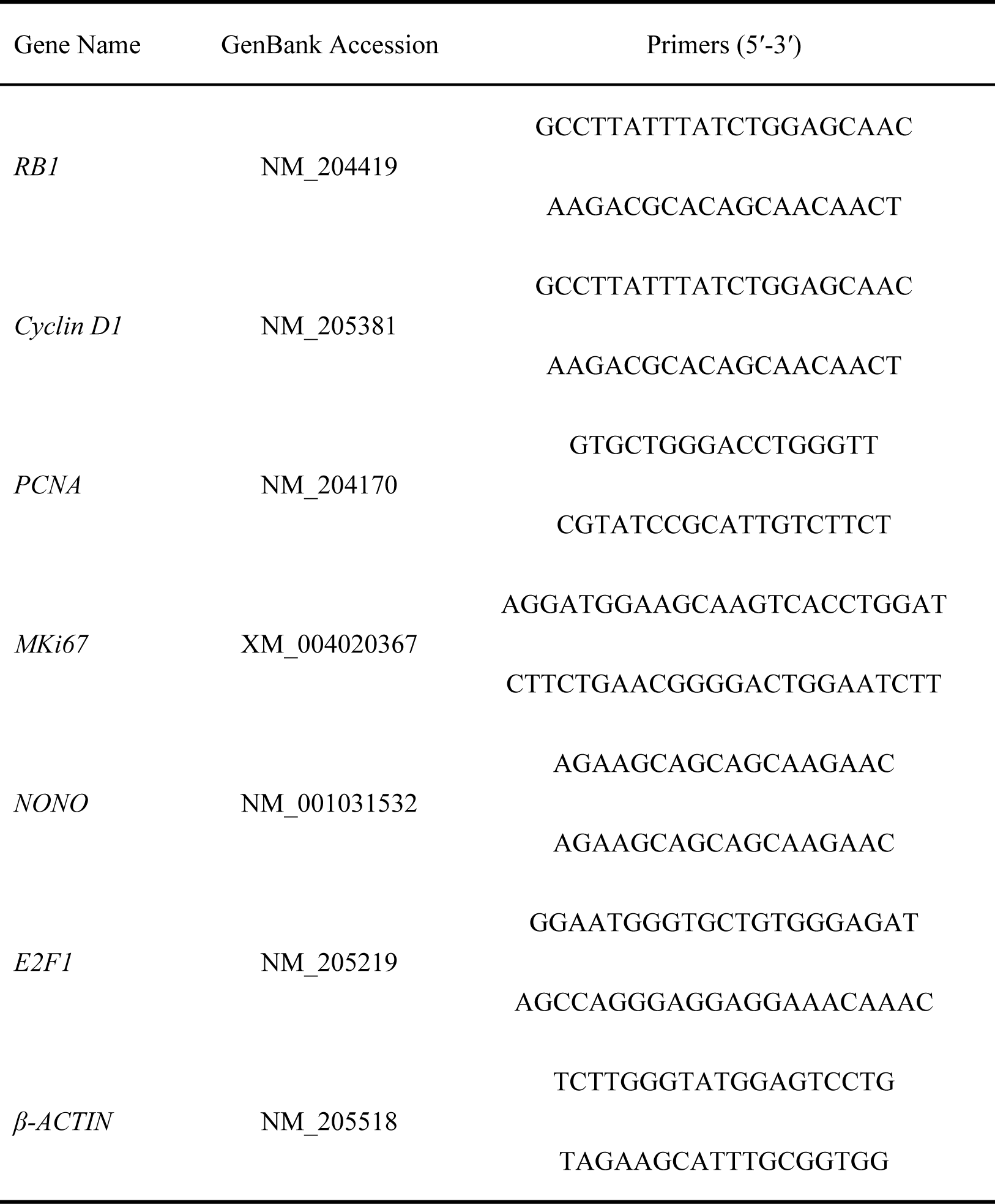
Primer sequences used for real-time RT-PCR.

### Western blot assays

Western blot analysis was used to examine the effect of *gRB1* overexpression. Chicken preadipocytes transfected with pCMA-HA-*PPARα* (M1, positive control), pCMV-HA-*gRB1* (M2, g*RB1* overexpression), or pCMV-HA (M3, control) vector for 2 days were homogenized in RIPA buffer (PBS, pH 7.4, containing 1% NP-40, 0.5% sodium deoxycholate, 0.1% SDS, and a protease inhibitor cocktail) supplemented with protease inhibitors (1 mmol/L phenylmethylsulfonyl fluoride, 0.002 g/L aprotinin, and 0.002 g/L leupeptin). Cellular debris and lipids were eliminated by centrifuging the solubilized samples at 13,000 × *g* for 60 min at 4°C. Cell lysates were separated by 5–12% SDS-polyacrylamide gels and transferred to polyvinylidene difluoride membranes. To block nonspecific binding, the membrane was incubated in blocking buffer (PBS with 5% nonfat dry milk) for 1 h at room temperature. After incubation with the primary antibody for HA-tag (1:200; Clontech) or β-ACTIN (1:1000; TransGen), a secondary horseradish peroxidase-conjugated antibody was added, and then a BeyoECL Plus kit (Beyotime, Jiang Su, China) was used for visualizing the protein bands.

### The 3-(4,5-dimethylthiazol-2-yl)-2,5-diphenyltetrazolium bromide (MTT) assay

The MTT assay was used to examine the effects of overexpression or knockdown of *gRB1* on ICP1 cells proliferation. ICP1 cells were transfected with the *gRB1*-overexpression plasmid or siRNA nucleotides for the knockdown of *RB1* expression, as well as the control plasmid or negative control nucleotides (siNC), respectively. After a 24-h incubation, these cells were passaged and seeded in 96-well plates at a concentration of 5000 cells per well. At the designated time points (including 12, 24, 48, 72, and 96 h), 20 mL of MTT solution (5 mg/mL; Sigma) were added to the medium, and the cells were incubated at 37°C for 4 h. After removal of the medium, 200 mL of DMSO were added to each well, and the plates were shaken on a rocking platform at 60 × *g* for 15 min. The cell solution was collected and the absorbance was recorded with an enzyme-labeled instrument (Bio-Rad, Hercules, CA, USA) at 492 nm. The experiments were repeated biologically three times. For each biological repeat, there were two groups, treatment group and control group, and cells were collected at 12, 24, 48, 72, and 96 h. At each time point, there were three wells technical duplications. Only the result of one biological repeat was shown in the current study, because the results of the three biological repeats were similar with each other.

### Cell-cycle analysis

After overexpression or knockdown of *gRB1* for 48 h, ICP1 cells were trypsinized and subsequently fixed with ice-cold 70% ethanol for at least 1 h. After extensive washing, the cells were suspended in propidium iodide (PI) staining solution (Beyotime), fully resuspended slowly, and bathed at 37°C for 30 min in the dark for subsequent FACScan analysis (Becton-Dickinson, San Jose, CA, USA). Cell-cycle analysis was performed by the ModFit LT software (Verity Software House, Topsham, ME, USA). The experiments were repeated biologically three times. Each biological repeat included three wells technical duplications. Only the result of one biological repeat was shown in the current study, because the results of the three biological repeats were similar with each other.

### Statistical analysis

Data are expressed as mean ± SD. Comparison between two groups was performed by the unpaired two-tailed Student’s t-test. Statistical analysis among more than two groups was performed using ANOVA method.

## Results

### gRB1 gene expression during chicken preadipocyte proliferation

The expression of *gRB1* during the proliferation of chicken preadipocytes was analyzed using real-time PCR. The cells were collected at 24, 48, 72 and 96 h, and the *NONO* gene was used as the internal reference. The results showed that the mRNA expression level of *gRB1* in chicken preadipocytes initially increased and then decreased, with a peak at 72 h (Fig 1).

**Fig 1.**
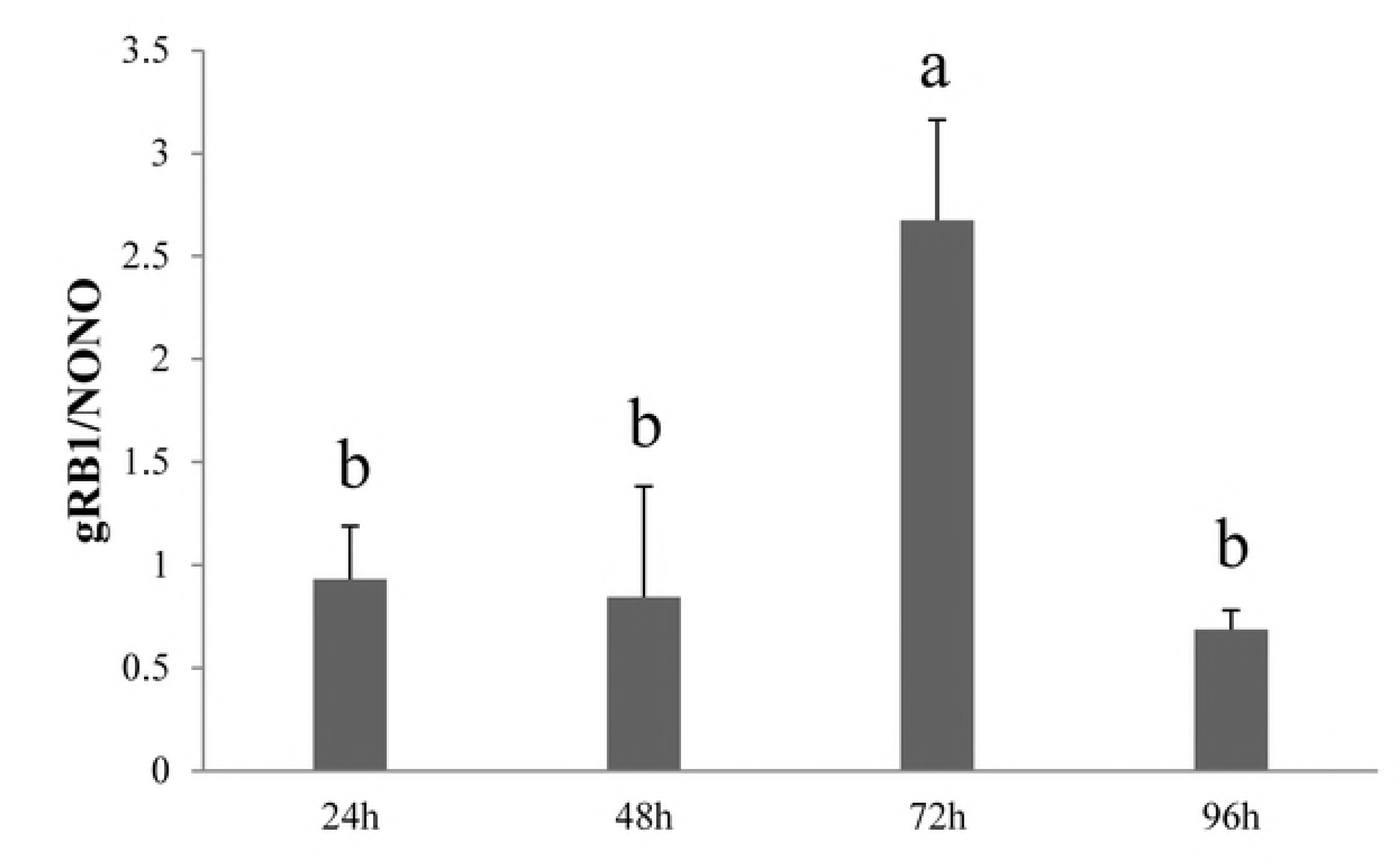
Expression of *gRB1* during chicken preadipocyte proliferation. The *gRB1* mRNA expression during chicken preadipocyte proliferation was detected by RT-qPCR. Cells were harvested at designated time points of 24, 48, 72 and 96 h. The *NONO* was used as the internal reference gene. Note: different lowercase letters above the columns indicate significant differences (*P* < 0.05).

### Effect of gRB1 on chicken preadipocyte proliferation

To explore the effect of *gRB1* on the proliferation of chicken preadipocytes, overexpression and RNAi experiments were carried out. The pCMV-HA-*PPARα* plasmid with a length of 52.19 kDa, which we successively constructed previously, was used as the positive control, and the pCMV-HA was used as the negative control. The western blot analysis results showed that pCMV-HA-*gRB1* expressed the RB1 protein in chicken preadipocytes (Fig 2A). The MTT assay results showed that overexpression of *gRB1* significantly suppressed the proliferation of ICP1 cells at 24 and 48 h (Fig 2B).

**Fig 2.**
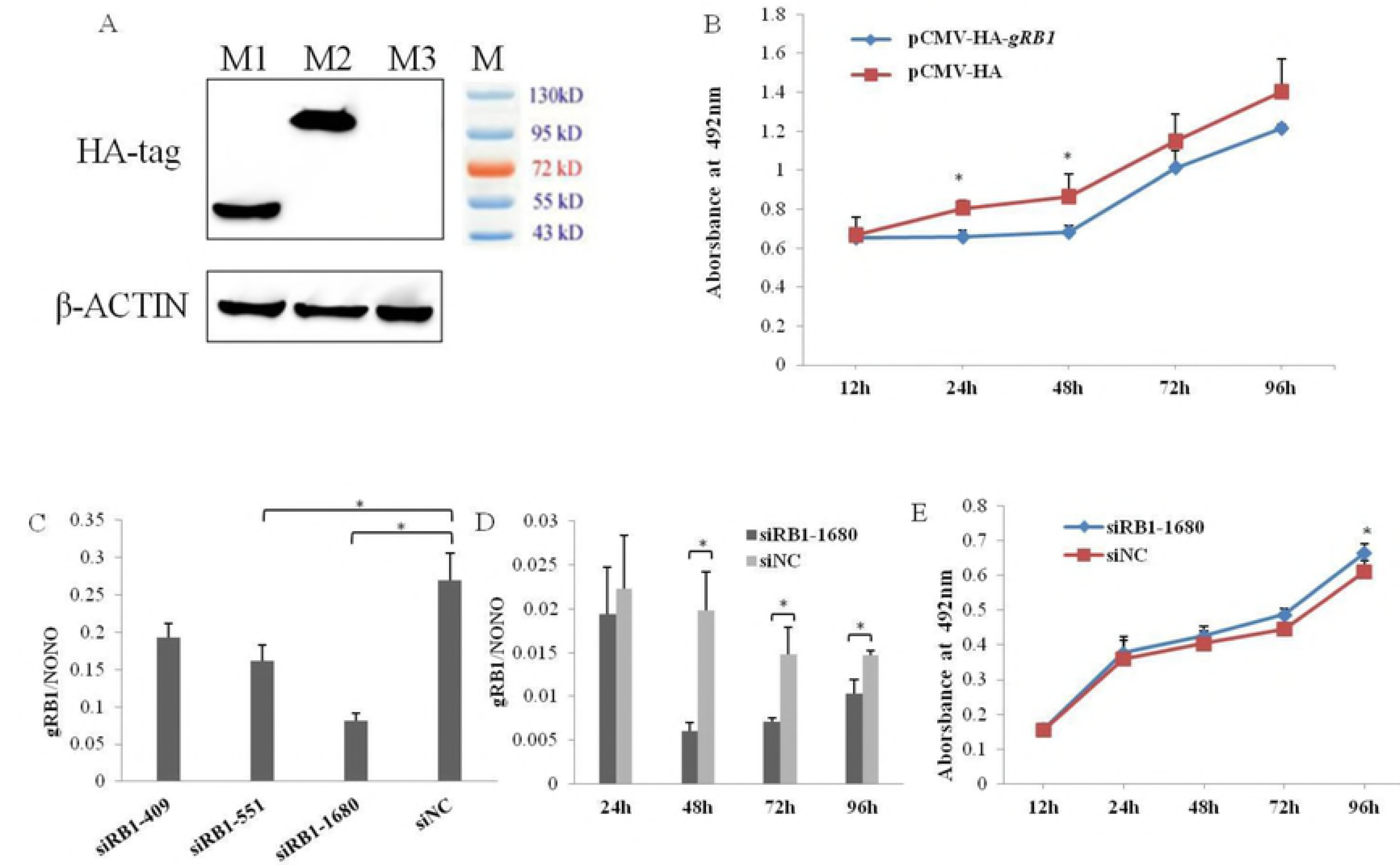
The analysis of the effect of g*RB1* gene on proliferation of chicken preadipocytes using overexpression and RNAi methods. (A) Validation of g*RB1* overexpression in chicken ICP1 cells by western blot. M1, pCMV-HA-*PPARα*, positive control, 52.19 kDa; M2, pCMV-HA-*gRB1*, g*RB1* overexpression, 104 kD; M3, pCMV-HA, EV; M, protein marker. (B) The analysis of the effect of g*RB1* overexpression on proliferation of chicken ICP1 cells by MTT assay. (C) Detection of the interference effect of different interference segments on *gRB1* expression in chicken ICP1 cells. (D) The interference effect of si*RB1*-1680 on the expression of *gRB1* in chicken ICP1 cells at different time points. (E) The analysis of the effect of g*RB1* knockdown on proliferation of chicken ICP1 cells by MTT assay. The diagrams show the proliferation of ICP1 cells measured by absorbance at 492 nm (MTT), error bars represent the standard deviation of three replicates. Note: * indicates significant difference (*P* < 0.05).

For the RNAi experiment three interference fragments were used, which were named si*RB1*-409, si*RB1*-551 and si*RB1*-1680. The interference effect of these three different siRNAs was examined and the results indicated that si*RB1*-551 and si*RB1*-1680 significantly knocked down the *gRB1* gene expression level compared with the control group (*P* < 0.05; Fig 2C). si*RB1*-1680 had the strongest interference effect, hence it was used to investigate the role of the *gRB1* gene in the proliferation of chicken preadipocytes in the following experiment. We analyzed the interference effect of si*RB1*-1680 on *gRB1* expression at different time points and found that the knockdown effect was the most significant at 48 h (Fig 2D). The MTT results indicated that knockdown of *gRB1* promoted the proliferation of ICP1 cells and that the effect was significant at 96 h compared with the negative control (Fig 2E).

### Effect of gRB1 on the cell cycle of chicken preadipocytes

The MTT results indicated that *gRB1* suppressed the proliferation of preadipocytes. To explore whether the suppression is caused by changes in the cell cycle, the effects of *gRB1* on the cell cycle of preadipocytes (ICP1) were analyzed using overexpression and RNAi methods. The effects of *gRB1* on the cell cycle of preadipocytes (ICP1) were examined at 48 h after transfection, because the overexpression and interference effects were the most significant at that time point. The overexpression results showed that the proportion of preadipocytes (ICP1) in the G1 and G2 phases significantly increased (*P* < 0.05), and the proportion of preadipocytes (ICP1) in the S phase significantly decreased (*P* < 0.05; Fig 3A). The RNAi results showed that the proportion of preadipocytes (ICP1) in the S phase significantly increased (*P* < 0.05) after the downregulation of the *gRB1* gene (Fig 3B).

**Fig 3.**
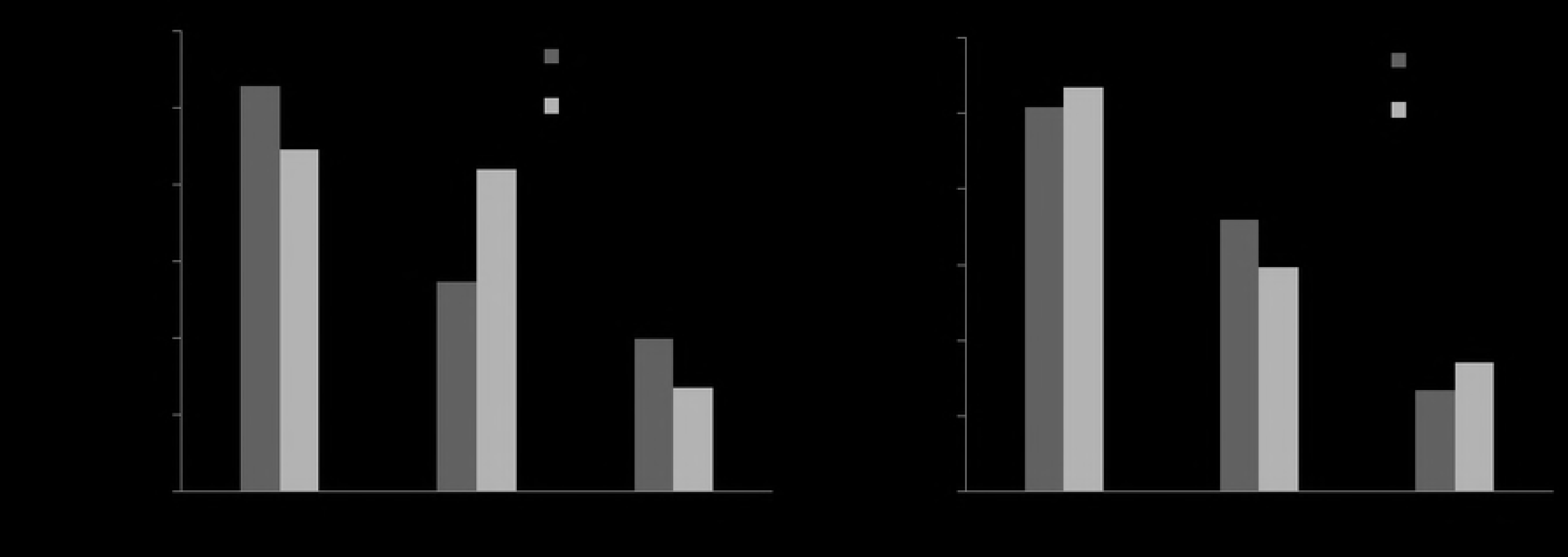
*gRB1* inhibits the G1/S transition of the cell cycle in ICP1 cells at 48 h after transfection. (A) The effects of *gRB1* overexpression on ICP1 cell cycle. (B) The effects of *gRB1* interference on ICP1 cell cycle. Note: * indicates significant difference (*P* < 0.05).

### Effect of gRB1 on the expression levels of genes related to chicken preadipocyte proliferation

In the current study, the effects of *gRB1* on the expression levels of genes related to the proliferation of chicken preadipocytes, including *MKi67*, *PCNA*, *E2F1* and *Cyclin D1*, were also analyzed. ICP1 cells were seeded in 6-well plates. The time point at which cell confluence reached 30–50% was the 0 h. At this time point, si*RB1*-1680, siNC, pCMV-HA-*gRB1*, or pCMV-HA was transfected into the preadipocytes. The preadipocytes were collected at 24, 48, 72, and 96 h, and RNA was extracted. The *NONO* gene was used as the internal reference. The results showed that the expression level of *MKi67* significantly decreased at 72 h (*P* < 0.05) and the expression levels of *E2F1*, *PCNA* and *Cyclin D1* did not significantly change after *gRB1* overexpression (Fig 4A). In contrast, the expression level of *MKi67* significantly increased at 96 h (*P* < 0.05), the expression levels of *E2F1*, *PCNA* and *Cyclin D1* did not significantly change after *gRB1* gene knockdown (Fig 4B).

**Fig 4.**
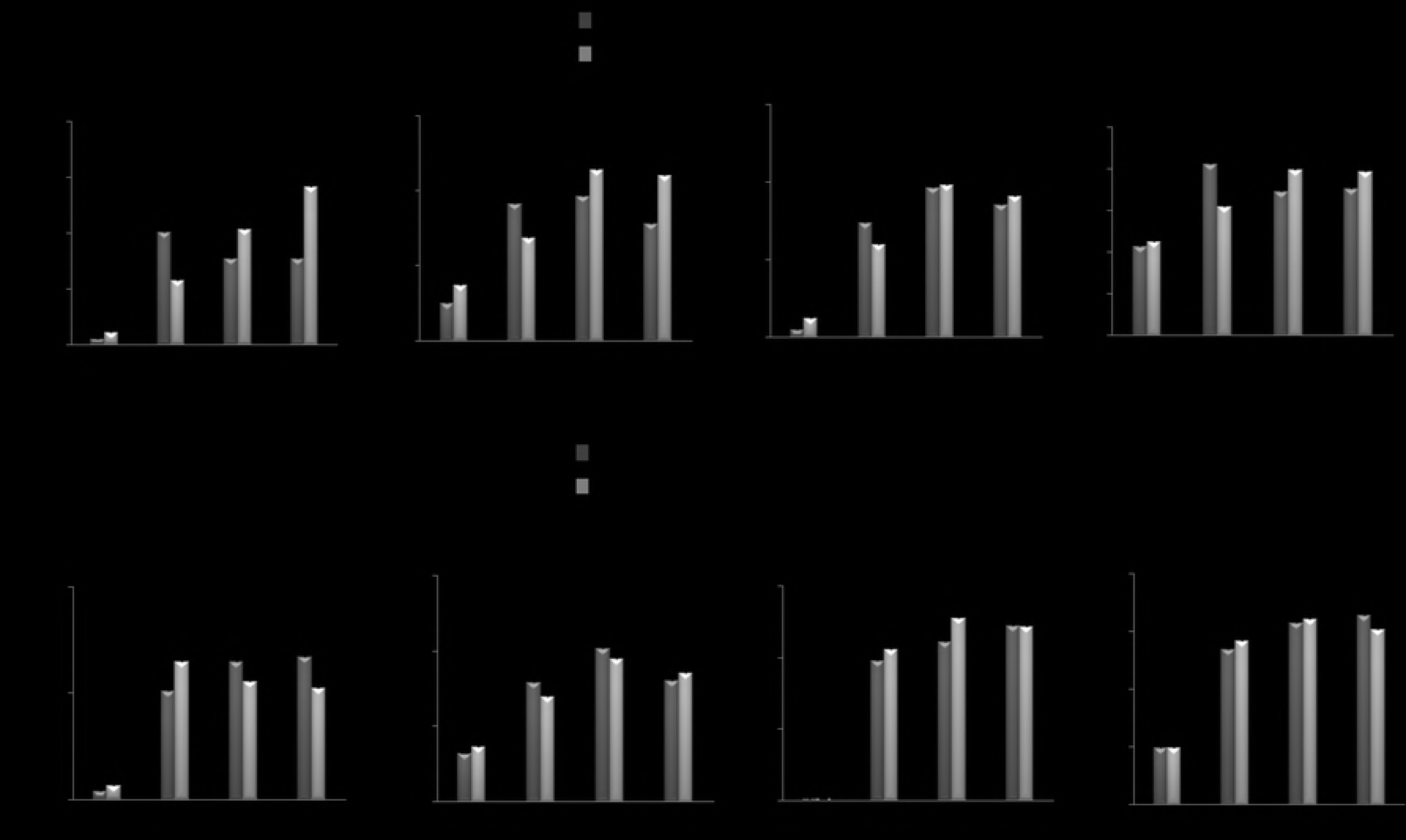
The effect of *gRB1* on the expression levels of genes related to the proliferation of ICP1 cells. (A) The effect of *gRB1* overexpression on the expression of important genes during ICP1 cell proliferation. (B) The effect of *gRB1* interference on the expression of important genes during ICP1 cell proliferation. Note: * indicates significant difference (*P* < 0.05).

## Discussion

In the current study, we used overexpression and RNAi methods to reveal the role of *gRB1* gene expression in the proliferation of chicken preadipocytes. The MTT results showed that overexpression of *gRB1* suppressed preadipocyte proliferation and *gRB1* knockdown promoted the proliferation of chicken preadipocytes, indicating that *gRB1* suppressed the proliferation of preadipocytes in chickens. These results are consistent with studies that showed that mouse *RB1* inhibited the proliferation of 3T3-L1 cells [9], knockdown of *RB1* in porcine could increase the mRNA levels of adipogenic markers, such as *PPARγ*, *aP2*, *LPL* and adiponectin [10], and human *RB1* inhibited the proliferation of cancer cells, including retinoblastoma [11], prostate cancer [12], and lung cancer cells [13].

Here, we found that the proportion of preadipocytes in the G1 phase significantly increased (*P* < 0.05) and the proportion of preadipocytes in the S phase significantly decreased (*P* < 0.05) after overexpression of the *gRB1*. In addition, when the expression level of *gRB1* gene was knocked down, the proportion of preadipocytes in the S phase significantly increased (*P* < 0.05) and the proportion of preadipocytes in the G1 phase had a tendency to decrease. These results indicated that *gRB1* inhibited the cell cycle of preadipocytes, mainly by inhibiting the G1 to S phase transition, resulting in G1 arrest. These results are consistent with studies on mammals that showed that RB1 is a negative regulator of the cell cycle, as it primarily binds to E2F, thereby preventing the progression of cells through the G1/S phase, resulting in cell-cycle arrest [14-17].

In our study, four cell-cycle-related genes, including *MKi67*, *PCNA*, *E2F1*, and *CyclinD1*, were selected to determine the mechanism of the effect of *gRB1* on the cell cycle. *MKi67* is a nuclear protein that is associated with and may be necessary for cellular proliferation [18,19]. Additionally, inactivation of *MKi67* leads to inhibition of ribosomal RNA synthesis [20,21]. The results of the current study showed that *gRB1* overexpression inhibited the expression level of *MKi67* at 72 h, and that *gRB1* knockdown promoted the expression level of *MKi67* at 96 h. These results indicated that *gRB1* decreased the expression level of *MKi67*. This result is consistent with studies showing that *RB1* suppressed the proliferation of human cells and that *MKi67* protein promoted cell proliferation [16,18].

Proliferating cell nuclear antigen (*PCNA*), a good indicator of cell proliferation, is a cofactor of DNA polymerase δ, and plays an important role in the initiation of cell proliferation [22-25]. The current study showed that the expression level of *PCNA* increased at 48 and 72 h after *gRB1* knockdown, and decreased at 72 and 96 h after *gRB1* overexpression using *NONO* as the internal reference gene. Though these results were not statistically significant, they imply that *gRB1* decreased the expression level of *PCNA*. This result is consistent with studies showing that *RB1* suppressed the proliferation of human cells and *PCNA* promoted cell proliferation [16,25].

*E2F1* acts as a transcription factor of genes involved in cell-cycle progression, DNA replication, DNA repair, and apoptosis [26,27]. In human cells, E2F1 binds preferentially to retinoblastoma protein (pRb) in a cell-cycle-dependent manner and can mediate both cell proliferation and p53-dependent/independent apoptosis [28]. Here, the mRNA expression level of *E2F1* did not significantly change after knockdown or overexpression of the *gRB1* gene. This suggests that the interaction between *gRB1* and *E2F1* may not be at the transcriptional level, but at the translation level, similarly to the case in mammals [28]. In mammals, pRb interacts with the E2F1 protein and plays important functions in DNA replication and cell proliferation [28,29].

Cyclin D1 is a cell-cycle protein and is an important regulatory factor of the G1 to S phase transition [30,31]. Cyclin D1 and E2F participate in the regulation of cell-cycle-based networks of pRb [32]. Here, the expression level of *Cyclin D1* did not significantly change after knockdown or overexpression of the *gRB1* gene. This result also indicates that the interaction between *gRB1* and *Cyclin D1* may not be at the transcriptional level, but at the translation level, as is the case in mammals [33,34]. In mammals, Cyclin D1 may be able to independently regulate the activity of pRb [34].

In summary, our results showed that *gRB1* is a negative regulator of chicken preadipocyte proliferation, and that it inhibited the cell cycle of preadipocytes through G1 arrest.

## Acknowledgement

This research was supported by the National Natural Science Foundation (No. 31301960), the National 863 Project of China (No. 2013AA102501), the China Postdoctoral Science Foundation (No. 2015M581421), the Heilongjiang Postdoctoral Financial Assistance (No. LBH-TZ0612), and the “Academic Backbone” Project of Northeast Agricultural University (No. 17XG18).

## References

1. Claire D’Andre H, Paul W, Shen X, Jia X, Zhang R, Sun L, et al. Identification and characterization of genes that control fat deposition in chickens. J Anim Sci Biotechnol. 2013; 4(1): 43.

2. de Verdal H, Narcy A, Bastianelli D, Même N, Urvoix S, Collin A, et al. Genetic variability of metabolic characteristics in chickens selected for their ability to digest wheat. J Anim Sci. 2013; 91(6): 2605–15.

3. Demeure O, Duclos MJ, Bacciu N, Le Mignon G, Filangi O, Pitel F, et al. Genome-wide interval mapping using SNPs identifies new QTL for growth, body composition and several physiological variables in an F2 intercross between fat and lean chicken lines. Genet Sel Evol. 2013; 45: 36.

4. Ramiah SK, Meng GY, Sheau Wei T, Swee Keong Y, Ebrahimi M. Dietary Conjugated Linoleic Acid Supplementation Leads to Downregulation of PPAR Transcription in Broiler Chickens and Reduction of Adipocyte Cellularity. PPAR Res. 2014; 2014: 137652.

5. Liu X, Li H, Wang S, Hu X, Gao Y, Wang Q, et al. Mapping quantitative trait loci affecting body weight and abdominal fat weight on chicken chromosome one. Poult Sci. 2007; 86(6): 1084–9.

6. Liu X, Zhang H, Li H, Li N, Zhang Y, Zhang Q, et al. Fine-mapping quantitative trait loci for body weight and abdominal fat traits: effects of marker density and sample size. Poult Sci. 2008; 87(7): 1314–9.

7. Zhang H, Wang SZ, Wang ZP, Da Y, Wang N, Hu XX, et al. A genome-wide scan of selective sweeps in two broiler chicken lines divergently selected for abdominal fat content. BMC Genomics. 2012; 13: 704.

8. Wang W, Zhang T, Wu C, Wang S, Wang Y, Li H, et al. Immortalization of chicken preadipocytes by retroviral transduction of chicken TERT and TR. PLoS One. 2017; 12(5): e0177348.

9. Shang W, Yang Y, Jiang B, Jin H, Zhou L, Liu S, et al. Ginsenoside Rb1 promotes adipogenesis in 3T3-L1 cells by enhancing PPARgamma2 and C/EBPalpha gene expression. Life Sci. 2007; 80(7): 618–25.

10. Hu X, Luo P, Peng X, Song T, Zhou Y, Wei H, et al. Molecular cloning, expression pattern analysis of porcine Rb1 gene and its regulatory roles during primary dedifferentiated fat cells adipogenic differentiation. Gen Comp Endocrinol. 2015; 214: 77–86.

11. Yang Y, Tian S, Brown B, Chen P, Hu H, Xia L, et al. The Rb1 gene inhibits the viability of retinoblastoma cells by regulating homologous recombination. Int J Mol Med. 2013; 32(1): 137–43.

12. Sharma A, Comstock CE, Knudsen ES, Cao KH, Hess-Wilson JK, Morey LM, et al. Retinoblastoma tumor suppressor status is a critical determinant of therapeutic response in prostate cancer cells. Cancer Res. 2007; 67(13): 6192–203.

13. Feng S, Cong S, Zhang X, Bao X, Wang W, Li H, et al. MicroRNA-192 targeting retinoblastoma 1 inhibits cell proliferation and induces cell apoptosis in lung cancer cells. Nucleic Acids Res. 2011; 39(15): 6669–78.

14. Krek W, Ewen ME, Shirodkar S, Arany Z, Kaelin WG Jr, Livingston DM. Negative regulation of the growth-promoting transcription factor E2F-1 by a stably bound cyclin A-dependent protein kinase. Cell. 1994; 78(1): 161–72.

15. Cho HS, Hayami S, Toyokawa G, Maejima K, Yamane Y, Suzuki T, et al. RB1 methylation by SMYD2 enhances cell cycle progression through an increase of RB1 phosphorylation. Neoplasia. 2012; 14(6): 476–86.

16. Zhang YF, Zhang AR, Zhang BC, Rao ZG, Gao JF, Lv MH, et al. MiR-26a regulates cell cycle and anoikis of human esophageal adenocarcinoma cells through Rb1-E2F1 signaling pathway. Mol Biol Rep. 2013; 40(2): 1711–20.

17. Lu Z, Bauzon F, Fu H, Cui J, Zhao H, Nakayama K, et al. Skp2 suppresses apoptosis in Rb1-deficient tumours by limiting E2F1 activity. Nat Commun. 2014; 5: 3463.

18. Scholzen T, Gerdes J. The Ki-67 protein: from the known and the unknown. J Cell Physiol. 2000; 182(3): 311–22.

19. Sánchez-Muñoz A, Plata-Fernández Y, Jaén A, Lomas M, Fernández M, Llácer C, et al. Proliferation determined by ki67 marker and pCR in locally advanced breast cancer patients treated with neo-adjuvant chemotherapy. Breast J. 2013; 19(6): 685–6.

20. Rahmanzadeh R, Hüttmann G, Gerdes J, Scholzen T. Chromophore-assisted light inactivation of pKi-67 leads to inhibition of ribosomal RNA synthesis. Cell Prolif. 2007; 40(3): 422–30.

21. Bando M, Iwakura H, Ariyasu H, Koyama H, Hosoda K, Adachi S, et al. Overexpression of intraislet ghrelin enhances β-cell proliferation after streptozotocin-induced β-cell injury in mice. Am J Physiol Endocrinol Metab. 2013; 305(1): E140–8.

22. Broich G, Lavezzi AM, Biondo B, Pignataro LD. PCNA--a cell proliferation marker in vocal cord cancer. Part II: Recurrence in malignant laryngeal lesions. In Vivo. 1996; 10(2): 175–8.

23. Chieffi P, Franco R, Fulgione D, Staibano S. PCNA in the testis of the frog, Rana esculenta: a molecular marker of the mitotic testicular epithelium proliferation. Gen Comp Endocrinol. 2000; 119(1): 11–6.

24. Bologna-Molina R, Mosqueda-Taylor A, Molina-Frechero N, Mori-Estevez AD, Sánchez-Acuña G. Comparison of the value of PCNA and Ki-67 as markers of cell proliferation in ameloblastic tumors. Med Oral Patol Oral Cir Bucal. 2013; 18(2): e174–9.

25. Zhao H, Chen MS, Lo YH, Waltz SE, Wang J, Ho PC, et al. The Ron receptor tyrosine kinase activates c-Abl to promote cell proliferation through tyrosine phosphorylation of PCNA in breast cancer. Oncogene. 2014; 33(11): 1429–37.

26. van den Heuvel S, Dyson NJ. Conserved functions of the pRB and E2F families. Nat Rev Mol Cell Biol. 2008; 9(9): 713–24.

27. Hallstrom TC, Nevins JR. Balancing the decision of cell proliferation and cell fate. Cell Cycle. 2009; 8(4): 532–5.

28. Sherr CJ, McCormick F. The RB and p53 pathways in cancer. Cancer Cell. 2002; 2(2): 103–12.

29. Nevins JR. The Rb/E2F pathway and cancer. Hum Mol Genet. 2001; 10(7): 699–703.

30. Coats S, Flanagan WM, Nourse J, Roberts JM. Requirement of p27Kip1 for restriction point control of the fibroblast cell cycle. Science. 1996; 272(5263): 877–80.

31. Shirali S, Aghaei M, Shabani M, Fathi M, Sohrabi M, Moeinifard M. Adenosine induces cell cycle arrest and apoptosis via cyclinD1/Cdk4 and Bcl-2/Bax pathways in human ovarian cancer cell line OVCAR-3. Tumour Biol. 2013; 34(2): 1085–95.

32. Ko LJ, Prives C. p53: puzzle and paradigm. Genes Dev. 1996; 10(9): 1054–72.

33. Dowdy SF, Hinds PW, Louie K, Reed SI, Arnold A, Weinberg RA. Physical interaction of the retinoblastoma protein with human D cyclins. Cell. 1993; 73(3): 499–511.

34. Siegert JL, Rushton JJ, Sellers WR, Kaelin WG Jr, Robbins PD. Cyclin D1 suppresses retinoblastoma protein-mediated inhibition of TAFII250 kinase activity. Oncogene. 2000; 19(50): 5703–11.

